# Comprehensive profiling of mutations to influenza virus PB2 that confer resistance to the cap-binding inhibitor pimodivir

**DOI:** 10.1101/2021.05.11.443605

**Authors:** Y.Q. Shirleen Soh, Keara D. Malone, Rachel T. Eguia, Jesse D. Bloom

## Abstract

Antivirals are used not only in current treatment of influenza, but are also stockpiled as a first line of defense against novel influenza strains for which vaccines have yet to be developed. Identifying drug resistance mutations can guide clinical deployment of the antiviral, and additionally define the mechanisms of drug action and drug resistance. Pimodivir is a first-in-class inhibitor of the polymerase basic protein 2 (PB2) subunit of the influenza A virus polymerase complex. A number of resistance mutations have previously been identified in treated patients or cell culture. Here, we generate a complete map of the effect of all single-amino-acid mutations to an avian PB2 on resistance to pimodivir. We identified both known and novel resistance mutations not only in the previously implicated cap-binding and mid-link domains, but also in the N-terminal domain. Our complete map of pimodivir resistance thus enables the evaluation of whether new viral strains contain mutations that will confer pimodivir resistance.

## 1. Introduction

Antivirals are an important prophylactic and treatment for influenza, particularly for high-risk patients and those with severe infections (1). They are also stockpiled as a first line of defense against pandemics caused by novel influenza strains (2), for which vaccines are unlikely to be available for many months.

The first influenza antivirals approved over two decades ago are the adamantanes, which target the M2 ion channel of influenza A viruses, and neuraminidase inhibitors (NAIs), which inhibit the viral enzyme that facilitates the release of viral progeny from infected cells (3). Their clinical use, however, has been impacted by emergence of resistance mutations. Consequently, adamantanes are no longer recommended for clinical use due to resistance in currently circulating influenza strains (4). NAIs remain the standard of care, though resistance has been shown to emerge in some patients (5), and went to fixation in the now extinct lineage of H1N1 influenza that circulated in humans prior to 2009 (6). As such, there is an important need for new antivirals that not only extend the clinically beneficial window of treatment, but also are effective against adamantane and NAI resistant strains.

Pimodivir (also known as VX-787) is a member of a new class of antivirals targeting the influenza polymerase complex (7). The influenza polymerase plays a pivotal role in replication and transcription of the viral genome (8, 9) and is highly conserved, and is thus an attractive antiviral target. Specifically, pimodivir targets the PB2 subunit of the polymerase, preventing its binding to the 7-methyl GTP caps of host-capped mRNAs, thus ultimately inhibiting the first step of viral gene transcription. Importantly, pimodivir is active against a diverse panel of influenza A virus strains, including H5N1 avian influenza (10), highlighting its potential as a first-line treatment against novel pandemic strains.

Identifying resistance mutations can guide clinical deployment of appropriate antivirals, and aid in developing new antivirals that are less susceptible to resistance. Known resistance mutations to pimodivir, as with previous influenza antivirals, have typically been identified one at a time as they arise over the course of antiviral treatment either in patients, or in the laboratory in cell culture (7, 10–12). These mutations are located primarily in the PB2 7-methyl GTP cap-binding pocket (10), as well as the mid-link domain where additional contacts with pimodivir are made (13, 14). However, these mutations likely do not represent the full range of mutations that affect pimodivir resistance.

Here, we fully map pimodivir resistance mutations in high-throughput. Specifically, using deep mutational scanning, we quantified how pimodivir resistance is affected by all amino-acid mutations to an avian influenza PB2 that are compatible with viral replication in human cells.

## 2. Materials and Methods

### 2.1 PB2 mutant virus libraries

We previously generated three independent mutant virus libraries containing all single amino-acid-mutations to PB2 from an avian influenza strain (15). The PB2 mutations were made on a background of reassortant virus using polymerase and nucleoprotein genes (PB2, PB1, PA, NP) from the avian influenza strain A/Green-winged Teal/Ohio/175/1986 (S009) (16), and remaining genes (HA, NA, M, NS) from A/WSN/1933 (H1N1) (17). These libraries were passaged in A549 cells to select for viruses that encode all functionally tolerated mutations in PB2 for viral replication in A549 cells. The resulting functional virus was used here for resistance profiling.

### 2.2. Resistance profiling

To identify resistance mutations, we passaged the mutant virus libraries in A549 cells in the absence or presence of pimodivir, and then identified the mutant viruses that were enriched upon drug selection using deep sequencing. We aimed to passage 1 × 10^6^ TCID50 of each mutant virus library in A549 cells at an MOI of 0.2 in the presence of 50 nM of pimodivir. 4 hours prior to infection, we seeded 5 × 10^6^ cells in D10 in a 15 cm dish. Just prior to the infection, we replaced D10 media with WGM with 50 nM of pimodivir. 1 × 10^6^ TCID50 of each virus library was then added to each plate. At 2 hours post-infection, we replaced the inoculum with fresh WGM with 50 nM pimodivir. 44 hour post-infection, we harvested the viral supernatant. As mock-selected controls, each mutant virus library was also similarly passaged in the absence of pimodivir. Selected and mock-selected viral supernatants were sequenced with a barcoded subamplicon sequencing approach as previously described (15, 18).

### 2.3. Analysis of deep sequencing data

Deep mutational scanning sequence data was analyzed using dms_tools2 (https://jbloomlab.github.io/dms_tools2, version 2.3.0). The differential selection (19) was quantified as the logarithm of the mutation’s enrichment in the pimodivir-selected mutant virus library relative to the mock-selected control library. Sequencing of wildtype DNA plasmid was used as the error control for calculating differential selection.

### 2.4. Data availability and source code

Code for the analyses is provided as Supplemental File 1 and at https://github.com/jbloomlab/PB2_Pimodivir_Resistance. Mean mutation and site differential selection measurements are provided as Supplemental File 2 and 3 respectively. Sequencing reads are deposited into the NCBI SRA under BioProject ID PRJNA719471.

### 2.5. Polymerase activity assays

Minigenome assays were performed in biological triplicate (starting from independent bacterial clones of each PB2 mutant) in HEK293T cells. We seeded 2.5 × 10^4^ HEK293T cells per well of a 96-well plate. Cells were transfected the next day with 10 ng each of HDM_S009_PB2 (for the respective mutant), HDM_S009_PB1, HDM_S009_PA, HDM_S009_NP, 30 ng of pHH-PB1-flank-eGFP reporter, and 30 ng of pcDNA-mCherry as transfection control using BioT. At 22 hr post-transfection, cells were trypsinized and analyzed by flow cytometry. We report minigenome activity as the percent of mCherry-positive cells that are GFP-positive.

## 3. Results

In a prior study, we generated triplicate mutant virus libraries containing all single-amino-acid mutations to PB2 from an avian influenza strain, S009, with the other polymerase complex genes (PB1, PA, and NP) also derived from S009, and the remaining genes derived from the lab-adapted A/WSN/1933 (H1N1) strain [15]. We passaged each library to obtain viruses capable of replication in the A549 human lung epithelial carcinoma cell line [15]. In the present study, we passaged these mutant virus libraries once more in A549 cells in the presence or absence of 50nM of pimodivir, a concentration chosen so that only a small fraction (1-10%) of viral titers was recovered compared to a mock selection (Fig S1A). To quantify the effect of each mutation on viral growth in the presence of pimodivir, we sequenced the passaged viruses, and then calculated the enrichment of each mutation in the pimodivir selected versus non-selected conditions. This value, termed hereafter the “differential selection”, reflects how favorable a mutation is in the presence over the absence of pimodivir.

Selection of pimodivir resistance mutations was highly reproducible across three biological replicates (Fig S1B). In addition to previously identified resistance mutations, we identified many more mutations, both at sites at which mutations were previously found, as well as new sites (Fig 1, Fig S2). Many of these mutations are evolutionarily accessible by single nucleotide substitutions from currently circulating PB2 sequences (Fig 1B-E). Resistance mutations occur in three regions of the PB2 protein.

**Figure 1.**
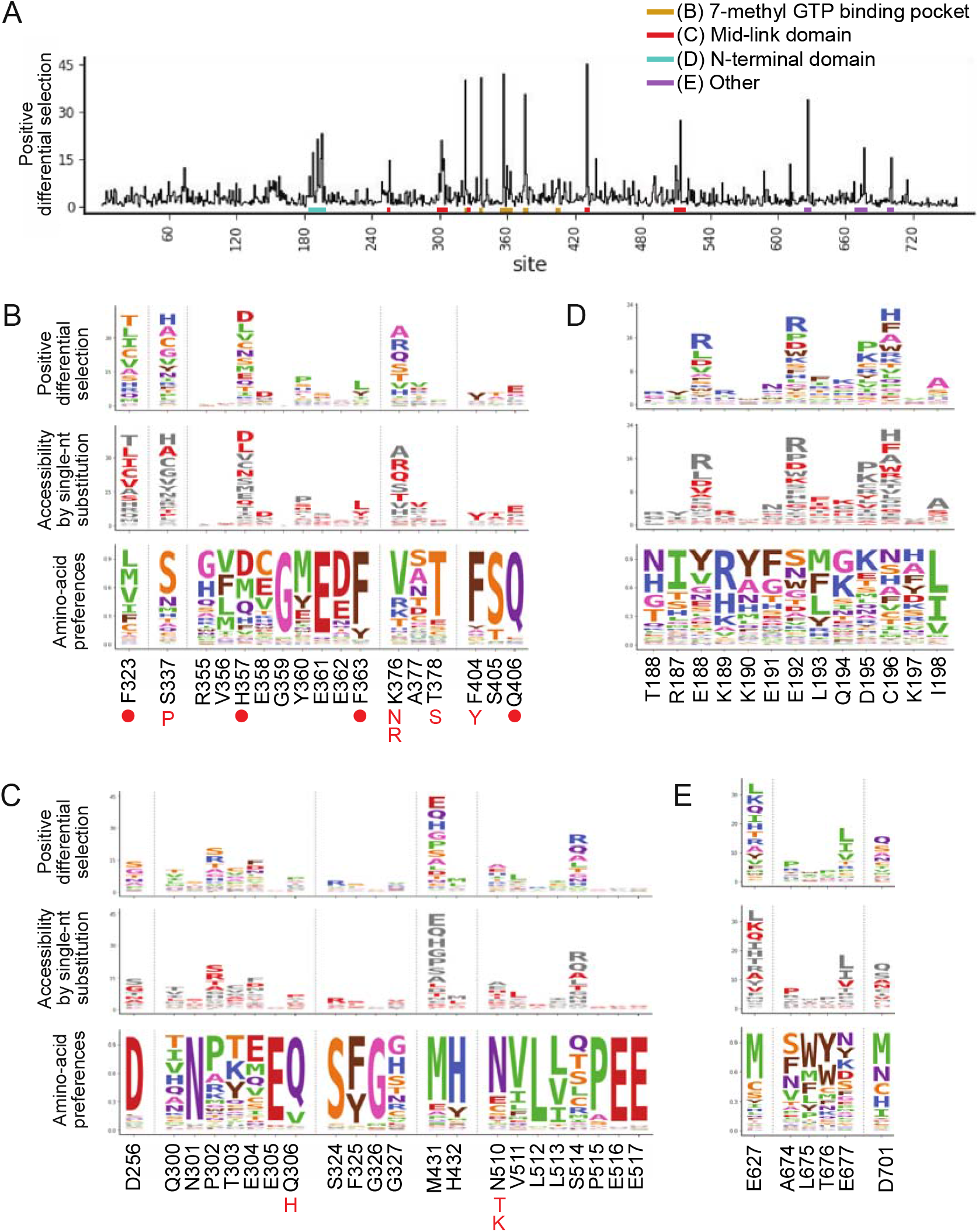
A complete map of pimodivir resistance. (A) Positive site differential selection across the entire PB2 protein. Regions of interest from different domains are underlined in different colors. (B—E) Mutation-level profiles for regions of interest in the (B) 7-methyl GTP binding pocket, (C) mid-link domain, (D) N-terminal domain, (E) other regions of PB2. The top plot of each panel shows the positive differential selection profile, where the height of each amino acid is proportional to its differential selection. The middle plot shows which amino acids are accessible by single-nucleotide substitution from existing avian PB2 sequences. Accessible amino acids are shown in red. The bottom plot shows the amino acid preferences at each site in human A549 cells, as previously measured in (15). The preference for an amino acid is proportional to its enrichment during the prior functional selection of the complete PB2 mutant library in A549 cells. The height of each letter is proportional to the preference for that amino acid at that site. Sites of known resistance mutations are indicated by either a circle underneath the site, or the specific resistance mutation if it is known.

The first region is the 7-methyl GTP cap-binding pocket (Fig 1B, 2A). Resistance mutations in this region, in combination with structural studies, provide support for the proposed mechanism that pimodivir occupies this pocket and interacts with PB2 similarly to the m7GTP guanine base [10]. Resistance mutations previously identified in this region are located at sites K376 which forms hydrogen bonds to both m7GTP and pimodivir, H357, F404, and F323 which form aromatic side chain interactions with the azaindole and pyrimidine rings of pimodivir, Q406 which interacts with the carboxylic acid of pimodivir, as well as sites S337, F363, and T378. For each of these sites with known resistance mutations, we identify additional substitutions at each of these sites that also confer resistance, e.g. sites 337 and 376. Sites that appear more tolerant of mutations, e.g. H357, have a correspondingly larger number of resistance mutations at that site, and vice versa, e.g. F404 (Fig 1B).

**Figure 2.**
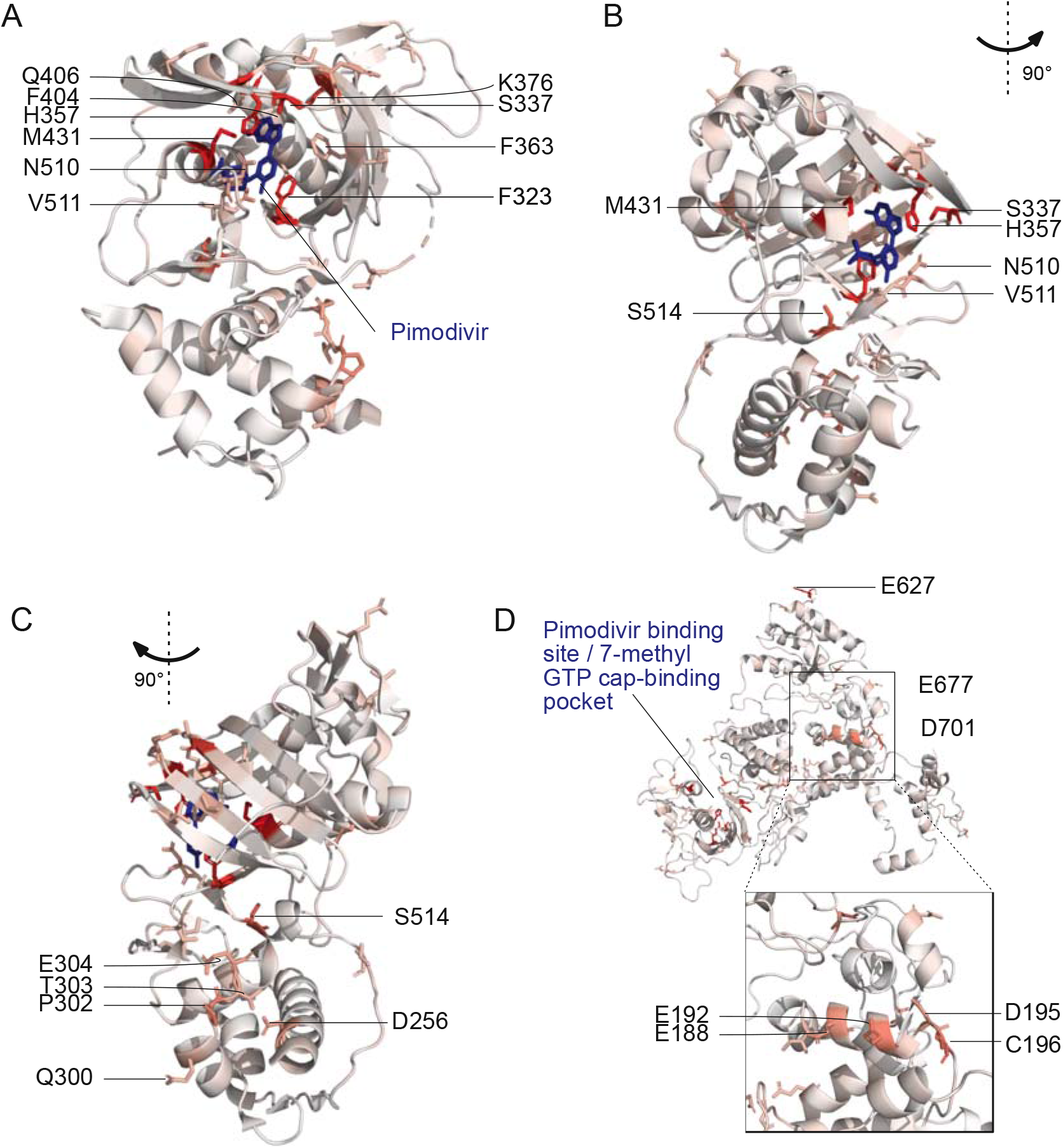
Locations of pimodivir resistance mutations on the PB2 protein. Amino acids are colored on a scale of white to red according to the total positive differential selection at that site, with red corresponding to high differential selection. Sites of interest, as identified in Figure 1, are labeled. (A-C) Structural model of PB2 cap-mid-link double domain with pimodivir in the transcriptionally inactive “apo” configuration [14]. Sites with resistance mutations that are in the 7-methyl GTP binding pocket and mid-link domain are labeled here. PDB: 6EUV (D) Structural model of full length PB2 in the apo configuration, but without pimodivir [20]. Sites of resistance mutations that are in the N-terminal domain and elsewhere are labeled here. PDB: 5D98

The second region is the mid-link domain (Figure 1C, 2B, 2C). Previous studies had identified resistance mutations, such as N510T [10], that lie outside the cap-binding pocket. More recent structural characterization of pimodivir in complex with the PB2 protein cap-binding and mid-link domain revealed that in the presence of pimodivir, the polymerase can also take up the transcriptionally inactive “apo” configuration. In this configuration, the mid-link and cap-binding domain are stabilized by inter-domain interactions, as well as by contacts between pimodivir and the mid-link domain [13,14]. Pimodivir binding is thus proposed to have a second indirect mode of action that is stabilization of this transcriptionally inactive “apo” state, thus occluding this cap-binding pocket. In support of this model, we identify numerous mutations at sites postulated to play a role in stabilizing the inhibitor bound “apo” state, such as at site

N510. We also identify resistance mutations at sites such as S514, which do not appear to directly contact pimodivir, but are in proximity to other segments of the mid-link domain. Substitution of serine by various bulky side-chains, such as arginine and glutamine, may destabilize the interaction between the cap-binding and mid-link domain, countering the stabilizing effect of pimodivir.

Unexpectedly, we identified resistance mutations in the PB2 N-terminal domain, such as at site E188, E192, D195, and C196, located far away from where pimodivir binds (Fig 1D, 2D). Strikingly, E188 and E195 are located side-by-side on an outward-facing side of an alpha-helix, and D195 and C196 are on an adjacent loop that is also on the surface of the PB2 protein. The most selected amino acids at these sites in the presence of pimodivir are basic amino acids including arginine, histidine, and lysine. Further investigation will be needed to fully understand the role of these sites in pimodivir resistance. Additionally, we also identified sites of high positive differential selection at sites known to be important in human-adaptation, namely 627 and 701 (Fig 1E). Thus, it appears that in the context of an avian influenza PB2, human adaptive mutations confer an advantage for influenza replication in human cells in the presence of pimodivir.

To validate that our high-throughput approach accurately identified mutations that counter the inhibitory effects of pimodivir on polymerase activity, we quantified the effect of mutations in a minigenome assay in the presence of a range of pimodivir concentrations (Fig 3A-C). We individually validated a selection of mutations from each of the three regions described above. A few new mutations were selected from two sites at which prior resistance mutations had been identified (F323L, H357D, M431E), and the remaining mutations are in sites that have not yet been associated with pimodivir resistance. Each mutation was made on the background of PB2 from the S009 avian influenza strain that has the E627K human-adaptive mutation. All mutations selected for validation increased pimodivir resistance, raising the EC50 between 2 to almost 30 fold (Fig 3E). Some of the resistance mutations, particularly those in the mid-link domain, appear to be detrimental to minigenome activity (Fig 3D). Hence, there may exist a trade-off between resistance to pimodivir, and baseline polymerase activity.

**Figure 3.**
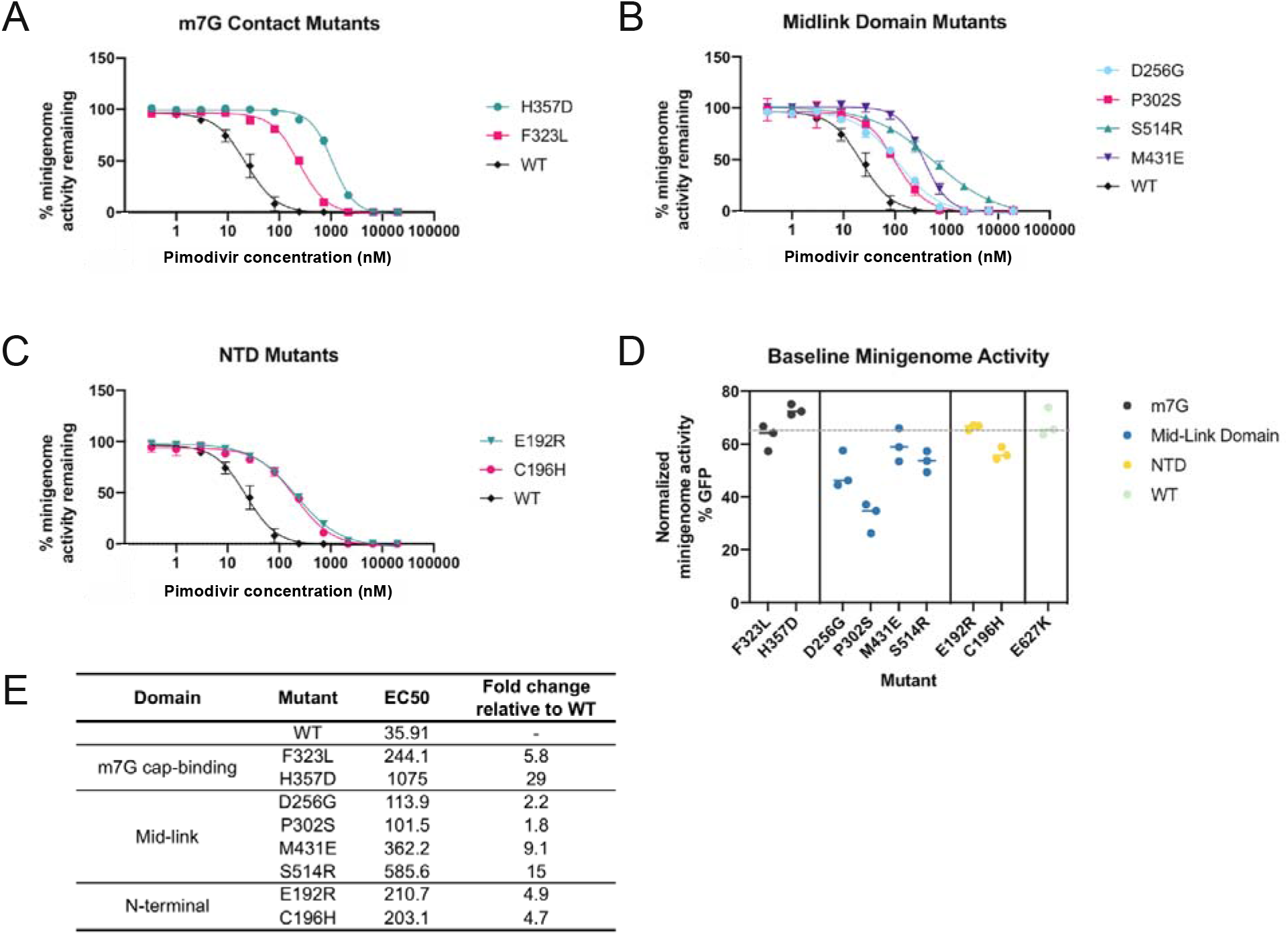
Validation of pimodivir resistance mutations using a minigenome assay. Mutations indicated were all made in the background of PB2-S009 with the E627K human-adaptive mutation. Hence, the “wildtype control”, “WT”, refers to PB2 that already has the E627K mutation. (A-C) Minigenome activity curves were plotted based on minigenome activity at various pimodivir concentrations, normalized to activity in the absence of pimodivir. (D) Minigenome activity was measured for polymerases containing each of the mutations, in the absence of pimodivir. (E) EC50 and fold-change in EC50 relative to wildtype, determined from the fit four-parameter logistic curves.

## 4. Discussion

We have quantified how pimodivir resistance is affected by all single-amino-acid mutations to an avian influenza PB2 that are compatible with viral replication in human cells. Our comprehensive mapping identified not only many previously known resistance mutations, but also new mutations at previously identified sites and new sites. The identified mutations in the cap-binding and mid-link regions of the PB2 protein lend support to current proposed mechanisms of drug action and thus resistance, namely that of direct inhibition of capped-RNA binding (10), as well as indirectly by stabilizing the transcriptionally inactive apo configuration of the cap-binding and mid-link domains (13, 14).

A third set of mutations in the PB2 N-terminal domain is unexplained by existing models of pimodivir action. Their characteristics suggest that they may interact with other domains or subunits of the polymerase or other proteins – based on existing structures they do not contact pimodivir directly. The site of these mutations (E188, E192, D195, C196) are in a tight cluster on the surface of the protein. Finally, basic amino acids appear to be favored across these sites. Further studies will be required to understand the role of these sites in pimodivir resistance. Finally, our comprehensive map of resistance mutations provides empirical data with which we can evaluate new viral strains for pimodivir resistance, including those mutations that would not have been predicted by structural analyses. This in turn will enable rapid determination of the best antiviral approach to be clinically deployed.

We began this work as pimodivir was in active development and in the midst of Phase 3 clinical trials. Recently, in September 2020, development of pimodivir for treatment of influenza was halted due to interim analyses of Phase 3 data that showed that pimodivir, while effective, did not demonstrate added benefit to current standard of care (https://www.janssen.com/janssen-discontinue-pimodivir-influenza-development-program). Our pimodivir resistance map, while not immediately applicable at the moment to an approved antiviral therapy, nevertheless is useful in case resistance against existing antivirals lead us to re-visit pimodivir as an alternate antiviral. Further, such maps of antiviral resistance may help us design the next generation of PB2 inhibitors that are less susceptible to evolution of resistance. More generally, we demonstrate the utility of deep mutational scanning to evaluate drug resistance, and thus facilitating informed clinical responses during pandemic outbreaks.

## Supplementary Materials

Figure S1: (A) The fraction of virus that survived pimodivir selection, as calculated by TCID50 recovered relative to the mock selected condition. (B) Correlations between the positive site differential selection for biological triplicates, Figure S2: Complete mutation-level resistance profile across PB2. The height of each amino acid is proportional to its differential selection, File S1: Code and results for computational analyses, File S2: Mean mutation differential selection measurements for pimodivir selection, File S3: Mean mutation and site differential selection measurements for pimodivir selection.

## Author Contributions

Conceptualization, Y.Q.S.S. and J.D.B.; investigation, Y.Q.S.S., K.D.M. and R.E.; resources, writing—original draft preparation, Y.Q.S.S.; writing—review and editing, Y.Q.S.S., K.D.M., R.E. and J.D.B. All authors have read and agreed to the published version of the manuscript.

## Funding

This research was supported by the NIAID of the NIH to J.D.B. (R01AI127893 and R01AI141707), Damon Runyon Cancer Research Foundation to Y.Q.S.S. (DRG-2271-16), and Mahan Postdoctoral Fellowship to Y.Q.S.S. J.D.B. is an Investigator of the Howard Hughes Medical Institute.

## Acknowledgments

We thank Jeffrey Taubenberger for providing plasmids for A/Green-winged Teal/Ohio/175/1986 (S009), and Andrew Mehle and Steven Baker for advice about growing virus containing S009 polymerase and NP genes. We thank William Fowler for help with molecular cloning.

## Conflicts of Interest

J.D.B. consults on viral evolution for Moderna, and has the potential to receive a share of PI revenue as an inventor on a Fred Hutch optioned technology/patent (application WO2020006494) related to deep mutational scanning of viral proteins. The funders had no role in the design of the study; in the collection, analyses, or interpretation of data; in the writing of the manuscript, or in the decision to publish the results

