## Supplementary figures and images for "Comprehensive profiling of mutations to influenza virus PB2 that confer resistance to the cap-binding inhibitor pimodivir"

### Figure S1

Figure S1

**A**

| Replicate | Fraction TCID50 recovered compared to mock selection |
| --- | --- |
| Rep1 | 1.0% |
| Rep2 | 10% |
| Rep3 | 1.0% |
| Wildtype | 0.56% |

**B**

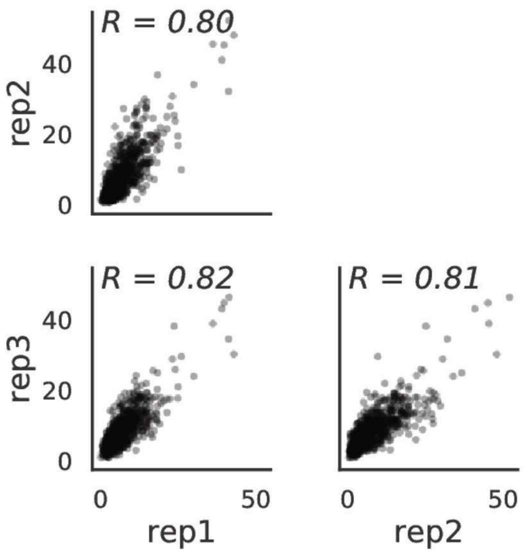

### Figure S2

#### Figure S2

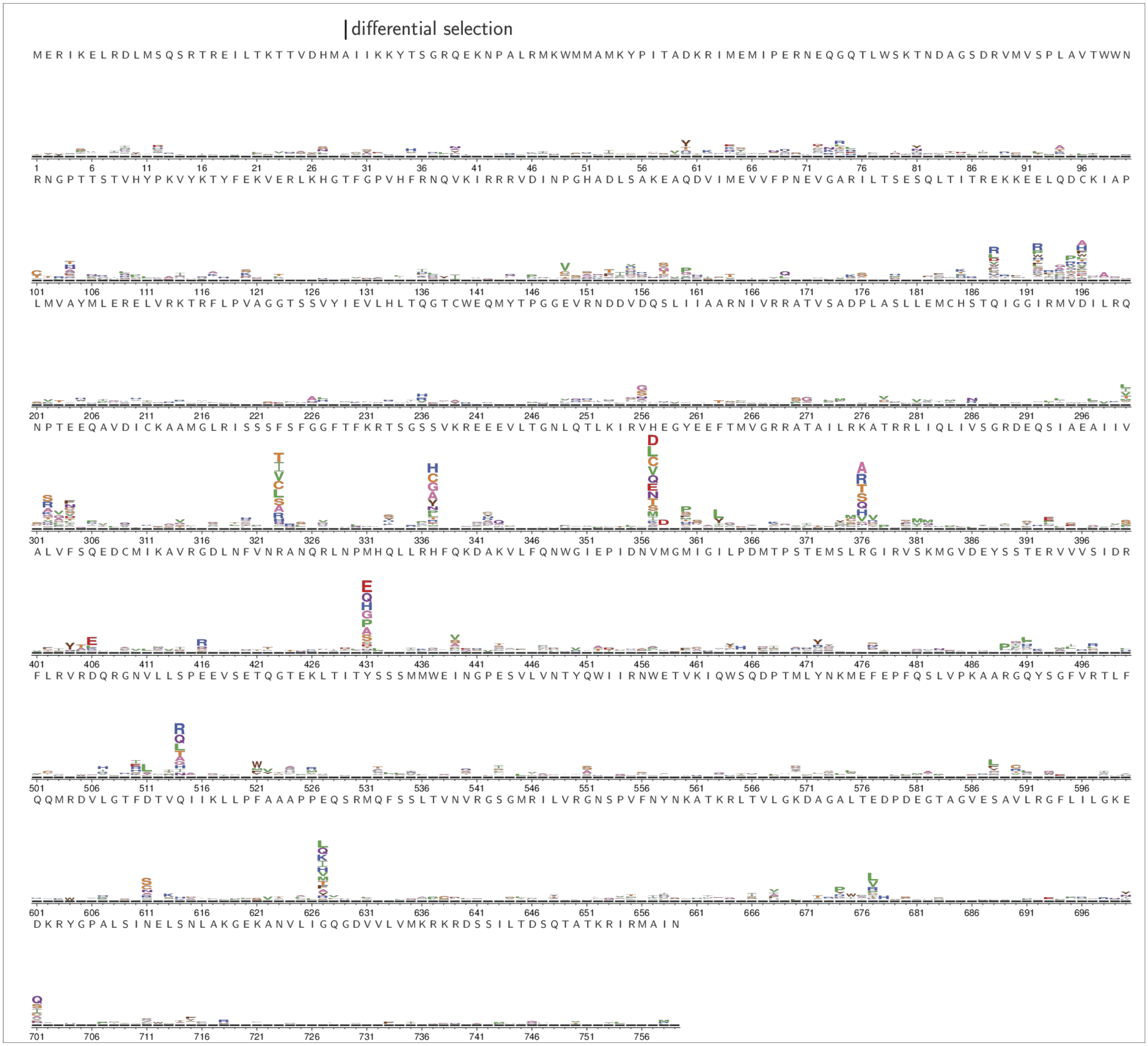
