## Supplementary material for "Comprehensive profiling of mutations to influenza virus PB2 that confer resistance to the cap-binding inhibitor pimodivir": File S1

analysis\_nb


### Comprehensive profiling of PB2 resistance mutations to pimodivir¶

#### Overview¶

Experiments and project led by Shirleen Soh, Keara Malone, and Rachel Eguia in the Bloom lab.

We comprehensively mapped PB2 resistance mutations to pimodivir, also known as VX-787. We use PB2 mutant virus libraries first described in Soh et al., 2019. The mutant libraries used here correspond to the three replicate libraries passaged on A549 cells in this paper.

Briefly, mutant virus libraries that were collected from previously passaging on A549 cells were passaged now on A549 cells in the presence or absence of pimodivir (`pimodivir-50nM`, or `mock`). The resulting viral variants were then deep sequenced to quantify the frequency of each mutation in each condition. Wildtype plasmid DNA was also sequenced to calculate the sequencing error rate.

We use dms\_tools2 to analyze the data. To identify mutations conferring resistance, we compute differential selection by comparing the frequency of mutations recovered after passaging in the presence versus absence of pimodivir.

#### Configure analysis¶

Import Python modules / packages.

We use dms\_tools2 for much of the analysis

In [58]:

```
import os
import subprocess
import warnings
import re

import numpy
import pandas as pd
from IPython.display import display, HTML
# import Bio.SeqIO

from Bio import SeqIO, AlignIO
from Bio.Data import CodonTable
from Bio.Seq import Seq


import matplotlib as mpl
import matplotlib.pyplot as plt
%matplotlib inline
import seaborn as sns

import dms_tools2
from dms_tools2.ipython_utils import showPDF
from dms_tools2.utils import codonEvolAccessibility
import dmslogo
from dmslogo.colorschemes import CBPALETTE

print(f"Using dms_tools2 version {dms_tools2.__version__}")

# CPUs to use, should not exceed the number you request with slurm
ncpus = 16

# do we use existing results or generate everything new?
use_existing = 'yes'


warnings.simplefilter('ignore')

# Set up directories
fastqdir = './fastq/'
datadir = './data/'
resultsdir = './results/' 
for xdir in [fastqdir, datadir, resultsdir]:
    if not os.path.isdir(xdir):
        os.mkdir(xdir)
resultsdir = {}  # holds name of directory for each result type
for result_type in ['codoncounts',
                    'diffsel'
                    ]:
    resultsdir[result_type] = os.path.join('results', result_type)
    os.makedirs(resultsdir[result_type], exist_ok=True)
    print(f"{result_type} go in {resultsdir[result_type]}")
```

```
Using dms_tools2 version 2.6.8
codoncounts go in results/codoncounts
diffsel go in results/diffsel
```

#### Read in sample information¶

Read in the list of samples and assign a unique name to each one:

In [2]:

```
# samples = pd.read_csv('samplelist.csv')
samples = (pd.read_csv('samplelist.csv')
           .assign(name=lambda x:
                   numpy.where(x['replicate'].isna(),
                               x['sample_condition'],
                               x['sample_condition'] + '-' + x['replicate']
                               )
                   )
           )

display(HTML(samples.to_html(index=False)))
SRAsamples = samples[['SRAprefix','run']].rename(
    columns={'SRAprefix':'name'})
display(HTML(SRAsamples.to_html(index=False)))
```

| sample\_condition | replicate | SRAprefix | run | name |
| --- | --- | --- | --- | --- |
| mutvirus-mock50 | rep1 | A549-Lib4-mock50 | SRR14139037 | mutvirus-mock50-rep1 |
| mutvirus-mock50 | rep2 | A549-Lib5-mock50 | SRR14139036 | mutvirus-mock50-rep2 |
| mutvirus-mock50 | rep3 | A549-Lib6-mock50 | SRR14139034 | mutvirus-mock50-rep3 |
| wtvirus-mock50 | NaN | A549-WT-mock50 | SRR14139035 | wtvirus-mock50 |
| mutvirus-pim-50nM | rep1 | A549-Lib4-pim-50nM | SRR14139041 | mutvirus-pim-50nM-rep1 |
| mutvirus-pim-50nM | rep2 | A549-Lib5-pim-50nM | SRR14139040 | mutvirus-pim-50nM-rep2 |
| mutvirus-pim-50nM | rep3 | A549-Lib6-pim-50nM | SRR14139039 | mutvirus-pim-50nM-rep3 |
| wtvirus-pim-50nM | NaN | A549-WT-pim-50nM | SRR14139038 | wtvirus-pim-50nM |
| DNA-WT | NaN | DNA-WT | SRR14139033 | DNA-WT |

| name | run |
| --- | --- |
| A549-Lib4-mock50 | SRR14139037 |
| A549-Lib5-mock50 | SRR14139036 |
| A549-Lib6-mock50 | SRR14139034 |
| A549-WT-mock50 | SRR14139035 |
| A549-Lib4-pim-50nM | SRR14139041 |
| A549-Lib5-pim-50nM | SRR14139040 |
| A549-Lib6-pim-50nM | SRR14139039 |
| A549-WT-pim-50nM | SRR14139038 |
| DNA-WT | SRR14139033 |

##### Obtain sequencing data¶

The deep mutational scanning FASTQ files are deposited on the Sequence Read Archive under the run numbers provided in the file data/SraRunTable\_DMS.txt. To download these files, we use the dms\_tools2.sra.fastqFromSRA function. This requires the fastq-dump and aspera programs to be installed at the specified paths. On the Hutch server, load `module load Aspera-Connect/3.9.6`.

In [6]:

```
print('Downloading FASTQ files from the SRA...')
dms_tools2.sra.fastqFromSRA(
    samples=SRAsamples,
    fastq_dump='fastq-dump', # valid path to this program on the Hutch server
    fastqdir=fastqdir,
    aspera=None,
    overwrite={'no':True, 'yes':False}[use_existing],
    passonly=True,
    no_downloads=False,
    ncpus=1
        )
print('Completed download of FASTQ files from the SRA')

print('Here are the names of the downloaded files now found in {0}'.format(fastqdir))
display(HTML(SRAsamples.to_html(index=False)))
```

```
Downloading FASTQ files from the SRA...
Completed download of FASTQ files from the SRA
Here are the names of the downloaded files now found in ./fastq/
```

| name | run | R1 | R2 |
| --- | --- | --- | --- |
| A549-Lib4-mock50 | SRR14139037 | A549-Lib4-mock50\_R1.fastq.gz | A549-Lib4-mock50\_R2.fastq.gz |
| A549-Lib5-mock50 | SRR14139036 | A549-Lib5-mock50\_R1.fastq.gz | A549-Lib5-mock50\_R2.fastq.gz |
| A549-Lib6-mock50 | SRR14139034 | A549-Lib6-mock50\_R1.fastq.gz | A549-Lib6-mock50\_R2.fastq.gz |
| A549-WT-mock50 | SRR14139035 | A549-WT-mock50\_R1.fastq.gz | A549-WT-mock50\_R2.fastq.gz |
| A549-Lib4-pim-50nM | SRR14139041 | A549-Lib4-pim-50nM\_R1.fastq.gz | A549-Lib4-pim-50nM\_R2.fastq.gz |
| A549-Lib5-pim-50nM | SRR14139040 | A549-Lib5-pim-50nM\_R1.fastq.gz | A549-Lib5-pim-50nM\_R2.fastq.gz |
| A549-Lib6-pim-50nM | SRR14139039 | A549-Lib6-pim-50nM\_R1.fastq.gz | A549-Lib6-pim-50nM\_R2.fastq.gz |
| A549-WT-pim-50nM | SRR14139038 | A549-WT-pim-50nM\_R1.fastq.gz | A549-WT-pim-50nM\_R2.fastq.gz |
| DNA-WT | SRR14139033 | DNA-WT\_R1.fastq.gz | DNA-WT\_R2.fastq.gz |

In [7]:

```
samples = samples.merge(SRAsamples[['run','R1']])
samples
```

Out[7]:

|  | sample\_condition | replicate | SRAprefix | run | name | R1 |
| --- | --- | --- | --- | --- | --- | --- |
| 0 | mutvirus-mock50 | rep1 | A549-Lib4-mock50 | SRR14139037 | mutvirus-mock50-rep1 | A549-Lib4-mock50\_R1.fastq.gz |
| 1 | mutvirus-mock50 | rep2 | A549-Lib5-mock50 | SRR14139036 | mutvirus-mock50-rep2 | A549-Lib5-mock50\_R1.fastq.gz |
| 2 | mutvirus-mock50 | rep3 | A549-Lib6-mock50 | SRR14139034 | mutvirus-mock50-rep3 | A549-Lib6-mock50\_R1.fastq.gz |
| 3 | wtvirus-mock50 | NaN | A549-WT-mock50 | SRR14139035 | wtvirus-mock50 | A549-WT-mock50\_R1.fastq.gz |
| 4 | mutvirus-pim-50nM | rep1 | A549-Lib4-pim-50nM | SRR14139041 | mutvirus-pim-50nM-rep1 | A549-Lib4-pim-50nM\_R1.fastq.gz |
| 5 | mutvirus-pim-50nM | rep2 | A549-Lib5-pim-50nM | SRR14139040 | mutvirus-pim-50nM-rep2 | A549-Lib5-pim-50nM\_R1.fastq.gz |
| 6 | mutvirus-pim-50nM | rep3 | A549-Lib6-pim-50nM | SRR14139039 | mutvirus-pim-50nM-rep3 | A549-Lib6-pim-50nM\_R1.fastq.gz |
| 7 | wtvirus-pim-50nM | NaN | A549-WT-pim-50nM | SRR14139038 | wtvirus-pim-50nM | A549-WT-pim-50nM\_R1.fastq.gz |
| 8 | DNA-WT | NaN | DNA-WT | SRR14139033 | DNA-WT | DNA-WT\_R1.fastq.gz |

##### Additional configuration¶

Specify the wildtype sequence for the gene:

In [8]:

```
refseqfile = './data/S009-PB2.fasta'

refseq = str(SeqIO.read(refseqfile, 'fasta').seq)

print(f"Read a reference sequence of {len(refseq)} nucleotides.")
```

```
Read a reference sequence of 2280 nucleotides.
```

#### Align sequencing and count mutations¶

The samples were sequenced using barcoded subamplicon sequencing to obtain high accuracy.
So we need to process these data to determine the counts of each codon mutation in each sample.

##### Run `dms2_batch_bcsubamp`¶

We process the seqeuncing data by using dms2\_batch\_bcsubamp to generate "codon counts" files that give the counts of each codon mutation for each sample.

###### Alignment parameters¶

Set alignment parameters used by dms2\_batch\_bcsubamp.
There is just a single subamplicon:

In [9]:

```
alignspecs = ' '.join(['1,285,36,32', 
                       '286,570,32,35', 
                       '571,855,37,32', 
                       '856,1140,32,35', 
                       '1141,1425,28,34', 
                       '1426,1710,33,34',
                       '1711,1995,29,29',
                       '1996,2280,28,43'])
r1trim = '200'
r2trim = '170'
```

###### Run the program¶

Write the batch file used as input by this program:

In [10]:

```
bcsubamp_batchfile = os.path.join(resultsdir['codoncounts'], 'batch.csv')

(samples
 [['name', 'R1']]
 .to_csv(bcsubamp_batchfile, index=False)
 )

print(f"Created batch file {bcsubamp_batchfile}")
```

```
Created batch file results/codoncounts/batch.csv
```

Now run the program:

In [13]:

```
cmds = ['dms2_batch_bcsubamp',
        '--batchfile', bcsubamp_batchfile,
        '--fastqdir', fastqdir,
        '--refseq', refseqfile,
        '--alignspecs', alignspecs,
        '--R1trim', r1trim,
        '--R2trim', r2trim,
        '--outdir', resultsdir['codoncounts'],
        '--summaryprefix', 'summary',
        '--ncpus', str(ncpus),
        '--use_existing', use_existing,
        '--bcinfo',  # save complete barcode info to look at subamplicons
        ]
cmd = ' '.join(cmds)

print(f"Running dms2_batch_bcsubamp with this command:\n{cmd}")
subprocess.check_output(cmds)
print('Completed running dms2_batch_bcsubamp.')
```

```
Running dms2_batch_bcsubamp with this command:
dms2_batch_bcsubamp --batchfile results/codoncounts/batch.csv --fastqdir ./fastq/ --refseq ./data/S009-PB2.fasta --alignspecs 1,285,36,32 286,570,32,35 571,855,37,32 856,1140,32,35 1141,1425,28,34 1426,1710,33,34 1711,1995,29,29 1996,2280,28,43 --R1trim 200 --R2trim 170 --outdir results/codoncounts --summaryprefix summary --ncpus 16 --use_existing yes --bcinfo
sbatch -c 16 --mem=300000 --time=24:00:00 --wrap="dms2_batch_bcsubamp --batchfile results/codoncounts/batch.csv --fastqdir ./fastq/ --refseq ./data/S009-PB2.fasta --alignspecs 1,285,36,32 286,570,32,35 571,855,37,32 856,1140,32,35 1141,1425,28,34 1426,1710,33,34 1711,1995,29,29 1996,2280,28,43 --R1trim 200 --R2trim 170 --outdir results/codoncounts --summaryprefix summary --ncpus 16 --use_existing yes --bcinfo"
Submitted batch job 66008080
```

##### Plot sequencing and mutation counts summaries¶

Running dms2\_batch\_bcsubamp created some summary plots.
The prefix on those plots is:

In [14]:

```
countsplotprefix = os.path.join(resultsdir['codoncounts'], 'summary_')
```

###### Reads and barcodes per sample¶

Total sequencing reads per sample:

In [15]:

```
showPDF(countsplotprefix + 'readstats.pdf')
```

Distribution of sequencing reads per barcode on subamplicons:

In [16]:

```
showPDF(countsplotprefix + 'readsperbc.pdf')
```

Number of barcoded subamplicons that align and have sufficient reads:

In [17]:

```
showPDF(countsplotprefix + 'bcstats.pdf')
```

Depth of valid barcoded subamplicons covering each site in the gene:

In [18]:

```
showPDF(countsplotprefix + 'depth.pdf')
```

###### Sampling of mutations¶

How thoroughly each mutation is sampled among the sequenced variants:

In [19]:

```
showPDF(countsplotprefix + 'cumulmutcounts.pdf')
```

Reassuringly, sampling generally drops in the pimodivir selected conditions compared to the mock selected.

###### Mutation frequencies¶

The average mutation frequency for each sample, stratifying by codon mutation type:

In [20]:

```
showPDF(countsplotprefix + 'codonmuttypes.pdf')
```

There appears to be selection for many non-synonymous mutations in the mutvirus libraries subject to pimodivir selection, relative to mock selected, as might be expected.

Average mutation frequency per sample, stratifying by number of nucleotide changes per codon mutation:

In [21]:

```
showPDF(countsplotprefix + 'codonntchanges.pdf')
```

Per-codon mutation frequencies across all sites in gene for each sample:

In [22]:

```
showPDF(countsplotprefix + 'mutfreq.pdf')
```

###### Check for oxidative damage¶

Sometimes there is oxidative damage which manifests as an enrichment of `G`->`T` and `C`->`A` mutations among the single-nucleotide codon mutations.
Check for this by plotting frequencies of different single-nucleotide mutation types:

In [23]:

```
showPDF(countsplotprefix + 'singlentchanges.pdf')
```

#### Analyze differential selection¶

We compute the differential selection on each mutation in paired sets of conditions.
This quantity indicates how much a mutation is favored in one condition versus the other.

##### Define the selection experiments¶

Each selection experiment compares two different conditions.

Here we compare replication in A549 cells in presence of pimodivir to no pimodivir. The 100nM and 50nM pimodivir selections are each paired with their respective mock selected samples.

Positive differential selection means a mutation is favored in the presence of pimodivir.

###### Define the comparisons¶

We define the names of the samples for the selected (*sel*) and mock (*mock*) sample for each selection type:

In [24]:

```
selectiontypes = pd.DataFrame(
                {'selection_type': ['Pimodivir50nM'],
                 'sel': ['mutvirus-pim-50nM'],
                 'mock': ['mutvirus-mock50'],
                 })
display(HTML(selectiontypes.to_html(index=False)))
```

| selection\_type | sel | mock |
| --- | --- | --- |
| Pimodivir50nM | mutvirus-pim-50nM | mutvirus-mock50 |

Now we make a data frame that gives the specific samples to compare for each selection, splitting them into groups based on the selection type.
For all samples, we use the *wildtype* sample as an error control (*err*):

In [31]:

```
selections = (
    selectiontypes
    .melt(id_vars='selection_type',
          value_vars=['sel', 'mock'],
          var_name='sample_type',
          value_name='sample',
          )
    .merge(samples, 
           left_on='sample', 
           right_on='sample_condition', 
           validate='one_to_many')
    .pivot_table(index=['selection_type', 'replicate'],
                 columns='sample_type',
                 values='name',
                 aggfunc=lambda x: ' '.join(x))
    .reset_index()
    .rename(columns={'selection_type': 'group',
                     'replicate': 'name'})
    .assign(err='DNA-WT')
    )
display(HTML(selections.to_html(index=False)))
```

| sample\_type | group | name | mock | sel | err |
| --- | --- | --- | --- | --- | --- |
|  | Pimodivir50nM | rep1 | mutvirus-mock50-rep1 | mutvirus-pim-50nM-rep1 | DNA-WT |
|  | Pimodivir50nM | rep2 | mutvirus-mock50-rep2 | mutvirus-pim-50nM-rep2 | DNA-WT |
|  | Pimodivir50nM | rep3 | mutvirus-mock50-rep3 | mutvirus-pim-50nM-rep3 | DNA-WT |

##### Run `dms2_batch_diffsel`¶

Now we run dms2\_batch\_diffsel to compute the differential selection.
Note that we can just use the *selections* data frame above as the batch file for input:

In [32]:

```
diffsel_batchfile = os.path.join(resultsdir['diffsel'], 'batch.csv')
selections.to_csv(diffsel_batchfile, index=False)

cmds = ['dms2_batch_diffsel',
        '--summaryprefix', 'summary',
        '--batchfile', diffsel_batchfile,
        '--outdir', resultsdir['diffsel'],
        '--indir', resultsdir['codoncounts'],
        '--use_existing', use_existing,
        '--ncpus', '1',
        ]
cmd = ' '.join(cmds)

print(f"Running dms2_batch_diffsel with this command:\n{cmd}")
subprocess.check_output(cmds)
print('Completed running dms2_batch_diffsel.')
```

```
Running dms2_batch_diffsel with this command:
dms2_batch_diffsel --summaryprefix summary --batchfile results/diffsel/batch.csv --outdir results/diffsel --indir results/codoncounts --use_existing yes --ncpus 1
sbatch -c 16 --mem=300000 --time=24:00:00 --wrap="dms2_batch_diffsel --summaryprefix summary --batchfile results/diffsel/batch.csv --outdir results/diffsel --indir results/codoncounts --use_existing yes --ncpus 1"
Submitted batch job 66103686
```

##### Plot site-level differential selection¶

We examine plots showing information about the differential selection.

First, we get the selection types to plot and the prefix for the summary out for each type:

In [33]:

```
diffsel_prefix = os.path.join(resultsdir['diffsel'], 'summary_')
seltype_prefixes = {s: f"{diffsel_prefix}{s}-"
                    for s in selectiontypes['selection_type']}
print('Here are the output prefixes:\t\n' +
      pd.Series(seltype_prefixes).to_csv(sep='\t')
      )
```

```
Here are the output prefixes:	
Pimodivir50nM	results/diffsel/summary_Pimodivir50nM-
```

###### Choose metrics to plot: absolute or positive differential selection¶

To summarize selection at a site level, we need to choose what metric to plot.
We have two options (see here for descriptions of both):

- *absolute site differential selection* , which includes the magnitude of selection for mutations that are favored in either of the two conditions being compared.
- \*positive site differential selection which only shows mutations favored in what we have called the **selected** condition.

###### Correlations among replicates¶

Correlation among the measurements for each selection type and replicate.

First for absolute site differential selection:

In [34]:

```
showPDF([f"{p}absolutesitediffselcorr.pdf" for p in seltype_prefixes.values()])
```

Now for positive site differential selection:

In [35]:

```
showPDF([f"{p}positivesitediffselcorr.pdf" for p in seltype_prefixes.values()])
```

###### Selection along the length of the gene¶

Now we plot the per-site selection across the gene, showing the *median* and *mean* of the libraries (replicates).
First for absolute differential selection.

In [36]:

```
showPDF(f"{diffsel_prefix}mediantotaldiffsel.pdf")
```

In [37]:

```
showPDF(f"{diffsel_prefix}meantotaldiffsel.pdf")
```

Now plot just the positive values:

In [38]:

```
showPDF(f"{diffsel_prefix}medianpositivediffsel.pdf")
```

In [39]:

```
showPDF(f"{diffsel_prefix}meanpositivediffsel.pdf")
```

##### Logo plots of selection on all mutations¶

We now use dms2\_logoplot to make logo plots showing the differential selection across all mutations.

In [40]:

```
for seltype, prefix in seltype_prefixes.items():
    
    for metric in ['all', 'positive']:

        diffselfile = f"{prefix}meanmutdiffsel.csv"
        logoplot = os.path.join(resultsdir['diffsel'],
                                f"{seltype}-{metric}_diffsel.pdf")

        print('\n\n' + ' '.join(['-' * 30, seltype, f"({metric} diffsel)",
                                 '-' * 30]) +
              '\n' + f"Plotting diffsel in {diffselfile} to {logoplot}\n")

        cmds = ['dms2_logoplot',
                '--diffsel', diffselfile,
                '--restrictdiffsel', metric,
                '--outdir', resultsdir['diffsel'],
                '--name', f"{seltype}-{metric}",
                '--nperline', '100', 
                '--numberevery', '5',
                '--scalebar', '10', 'differential selection',
                '--overlay1',  diffselfile, 'wildtype', 'wildtype',
                '--underlay', 'no',
                '--use_existing', use_existing,
                ]
        subprocess.check_output(cmds)

        showPDF(logoplot)
```

```
------------------------------ Pimodivir50nM (all diffsel) ------------------------------
Plotting diffsel in results/diffsel/summary_Pimodivir50nM-meanmutdiffsel.csv to results/diffsel/Pimodivir50nM-all_diffsel.pdf
```

```
------------------------------ Pimodivir50nM (positive diffsel) ------------------------------
Plotting diffsel in results/diffsel/summary_Pimodivir50nM-meanmutdiffsel.csv to results/diffsel/Pimodivir50nM-positive_diffsel.pdf
```

###### Interpretation and further analysis of differential selection logo plots¶

There appear to be only a handful of sites with very strong positive differential selection.

#### Select mutants for validation¶

I identified sites that were differentially selected in the presence of pimodivir. Based on prior literature and mapping onto the polymerase structure, the sites fell into roughly three regions:

1. Sites likely to be in direct contact with pimodivir or the m7G of mRNAs. These sites are fairly well-described in literature, though the few specific escape mutations have been identified even for sites thought to be important. Our experiments identify many escape mutations at these sites. I pick 2 mutations to validate here.
2. Sites that do not directly contact pimodivir/m7G, but rather, in the presence of pimodivir, stabilizes the apo-form of the polymerase, that excludes active transcription/replication. These were more recently described in the literature. Our experiments identify additional sites close to those sites that were previously thought to be implicated in this function. I pick 4 mutations to validate (2 each from slightly different locations), all from sites previously not implicated.
3. A cluster of sites in the N-terminal domain. Not previously described in literature. I wonder if it’s an artefact of our experiment. I pick 2 here to validate.

(I also noted high differential selection at sites I previously identified in my human vs duck cells selection. Hence, I suspect that mutations at these are more likely non-specifically improving polymerase function in human cells. (And I rationalize that this improvement in function is more evident in the presence of an inhibitor.) I did not pick any to validate here.)

I shortlisted the mutatons to test based on the following criteria:

1. At sites with high site differential selection
2. High mutation differential selection
3. Consistent across all 3 replicate libraries

##### Collate all data into single table for querying.¶

First, I collate site data for differential selection in pimodivir resistance, differential selection in human vs duck cells, and append information on whether it is a site already known to be involved in pimodivir resistance. Half of the top 10 sites are known, and half are new.

In [41]:

```
# site differential selection
allsitediffselmean = (pd.merge(pd.read_csv("results/diffsel/summary_Pimodivir50nM-meansitediffsel.csv"), 
                                pd.read_csv("data/summary_A549vCCL141-meansitediffsel.csv"), 
                               on='site', suffixes=('_pim','_A549'))
                       )

knownSites = set(pd.read_table("data/knownResistance.txt")['Site'])
allsitediffselmean["known"] = numpy.where(allsitediffselmean["site"].isin(knownSites), "yes", "no")
allsitediffselmean = allsitediffselmean[['site', 'abs_diffsel_pim', 'positive_diffsel_pim',
                                             'abs_diffsel_A549', 'positive_diffsel_A549',
                                             'known'
                                            ]]
allsitediffselmean.sort_values('positive_diffsel_pim', ascending=False).head(n=10)
```

Out[41]:

|  | site | abs\_diffsel\_pim | positive\_diffsel\_pim | abs\_diffsel\_A549 | positive\_diffsel\_A549 | known |
| --- | --- | --- | --- | --- | --- | --- |
| 0 | 431 | 44.999962 | 44.934134 | 29.827025 | 0.518602 | yes |
| 1 | 357 | 41.789181 | 41.789181 | 17.474837 | 14.032147 | yes |
| 2 | 337 | 40.598895 | 40.598895 | 10.303483 | 1.700791 | yes |
| 3 | 323 | 39.914311 | 39.876937 | 21.051950 | 19.962100 | yes |
| 5 | 376 | 35.435920 | 35.394002 | 12.082695 | 11.358961 | yes |
| 4 | 627 | 35.990894 | 33.737983 | 19.589407 | 18.009501 | no |
| 7 | 514 | 27.157629 | 27.157629 | 10.452031 | 5.027741 | no |
| 8 | 196 | 23.368766 | 23.030144 | 18.903253 | 0.198914 | no |
| 10 | 192 | 21.268584 | 21.257035 | 9.903482 | 6.898806 | no |
| 11 | 302 | 21.014377 | 20.913276 | 19.311978 | 0.439029 | no |

Next, I collate mutation data for differential selection in pimodivir resistance, differential selection in human vs duck cells, and append information on whether it is a site already known to be involved in pimodivir resistance. Almost all of the top 10 more differentially selected mutations in the presence of pimodivir are new!

In [45]:

```
# mutation differential selection
allmutdiffselmean = (pd.merge(pd.read_csv('results/diffsel/summary_Pimodivir50nM-meanmutdiffsel.csv'), 
                                pd.read_csv('data/summary_A549vCCL141-meanmutdiffsel.csv'),
                               on=['site','wildtype','mutation'], suffixes=('_pim','_A549')
                             ))
allmutdiffselmean["mut"] = (allmutdiffselmean["wildtype"] + 
                              allmutdiffselmean["site"].map(str) + 
                              allmutdiffselmean["mutation"])
knownMutations = set(pd.read_table("data/knownResistance.txt")['Mutation'].dropna())
allmutdiffselmean["known"] = numpy.where(allmutdiffselmean["mut"].isin(knownMutations), "yes", "no")
allmutdiffselmean.sort_values('mutdiffsel_pim', ascending=False, inplace=True)
allmutdiffselmean.head(n=10)
```

Out[45]:

|  | site | wildtype | mutation | mutdiffsel\_pim | mutdiffsel\_A549 | mut | known |
| --- | --- | --- | --- | --- | --- | --- | --- |
| 0 | 431 | M | E | 6.299282 | -1.367148 | M431E | no |
| 1 | 376 | K | A | 6.001965 | 0.834899 | K376A | no |
| 2 | 357 | H | D | 5.569827 | 2.337165 | H357D | no |
| 3 | 376 | K | R | 5.551795 | 2.105670 | K376R | yes |
| 4 | 337 | S | H | 5.468044 | 1.152390 | S337H | no |
| 5 | 677 | E | L | 5.416295 | -0.178586 | E677L | no |
| 6 | 514 | S | R | 5.367328 | 0.954636 | S514R | no |
| 7 | 323 | F | T | 5.364631 | 1.333052 | F323T | no |
| 8 | 431 | M | Q | 5.307767 | -0.514680 | M431Q | no |
| 9 | 376 | K | Q | 5.209439 | 0.804407 | K376Q | no |

Calculate the average minimum number of nucleotide substitutions needed to achieve a particular mutation.

In [46]:

```
def accessibility_threshold(row):
    if row['min Subst']==0:
        return '0'
    elif row['min Subst']<=1.1: # Picked this cut off to accommodate values slightly >1 when considering many sequences
        return '1'
    else:
        return '>1'
def seqsToCodonAcc(filename, seqlen):
    allowed_chars = set('ATCG')
    seqs = []
    for seq_record in SeqIO.parse(filename, 'fasta'):
        if len(seq_record.seq)==seqlen:
            if set(seq_record.seq).issubset(allowed_chars):
                seqs.append(str(seq_record.seq))
    #         else:
    #             print('Invalid bases:', seq_record.id, len(seq_record.seq))
#         else:
#             print('Not full length:', seq_record.id, len(seq_record.seq))
    accessibilitydf = (codonEvolAccessibility(seqs)
                     .melt(id_vars='site', value_vars=dms_tools2.AAS_WITHSTOP, 
                           var_name='toAA', value_name='min Subst')
                       .sort_values('site')
                    )
    accessibilitydf['acc'] = accessibilitydf.apply(lambda row: accessibility_threshold(row), axis=1)
    return accessibilitydf

accessibilityAvianPB2 = seqsToCodonAcc('data/Avian-PB2_2015-2019.fa', 2280)
accessibilityHumanPB2 = seqsToCodonAcc('data/Human-PB2_2019.fa', 2280)

accessibilitydf = pd.merge(accessibilityAvianPB2, accessibilityHumanPB2, how='inner',
                           on=['site', 'toAA'], suffixes=('_Avian', '_Human')
                          )
allmutdiffselmean = pd.merge(allmutdiffselmean, accessibilitydf, how='left',
                          left_on=['site', 'mutation'], right_on=['site', 'toAA'])
allmutdiffselmean.drop(columns=['mut', 'toAA'], inplace=True)
# Add columns for color of mutation, for logoplots to show accessibility
allmutdiffselmean['col_Avian'] = allmutdiffselmean['acc_Avian'].apply(lambda x: 'red' if x is '1' else 'gray')
allmutdiffselmean['col_Human'] = allmutdiffselmean['acc_Human'].apply(lambda x: 'red' if x is '1' else 'gray')
allmutdiffselmean["site_label"] = (allmutdiffselmean["wildtype"] + allmutdiffselmean["site"].map(str))
```

As expected, almost all the mutations known in the literature, which have been identified from patients or passaging in cell culture, are accessible by 1 nucleotide substitution.

In [47]:

```
allmutdiffselmean[allmutdiffselmean['known']=='yes']
```

Out[47]:

|  | site | wildtype | mutation | mutdiffsel\_pim | mutdiffsel\_A549 | known | min Subst\_Avian | acc\_Avian | min Subst\_Human | acc\_Human | col\_Avian | col\_Human | site\_label |
| --- | --- | --- | --- | --- | --- | --- | --- | --- | --- | --- | --- | --- | --- |
| 3 | 376 | K | R | 5.551795 | 2.105670 | yes | 0.999159 | 1 | 1.000000 | 1 | red | red | K376 |
| 32 | 404 | F | Y | 3.739456 | 0.864325 | yes | 1.000000 | 1 | 1.000000 | 1 | red | red | F404 |
| 62 | 324 | S | R | 2.935735 | -0.360399 | yes | 1.000000 | 1 | 1.000000 | 1 | red | red | S324 |
| 253 | 337 | S | P | 1.571473 | -0.755690 | yes | 1.000000 | 1 | 1.000000 | 1 | red | red | S337 |
| 320 | 510 | N | T | 1.402408 | -1.573899 | yes | 1.000841 | 1 | 1.000000 | 1 | red | red | N510 |
| 677 | 325 | F | L | 0.911180 | 0.072944 | yes | 1.000000 | 1 | 1.000245 | 1 | red | red | F325 |
| 698 | 510 | N | K | 0.890663 | -1.199185 | yes | 1.000560 | 1 | 1.000736 | 1 | red | red | N510 |
| 1004 | 376 | K | N | 0.688289 | 0.137758 | yes | 1.000841 | 1 | 1.000000 | 1 | red | red | K376 |
| 2374 | 306 | Q | H | 0.290695 | -0.339939 | yes | 1.000000 | 1 | 1.000000 | 1 | red | red | Q306 |
| 3176 | 324 | S | N | 0.174482 | -0.328176 | yes | 1.000000 | 1 | 1.000000 | 1 | red | red | S324 |
| 3518 | 378 | T | S | 0.145480 | -1.301878 | yes | 0.999440 | 1 | 1.000000 | 1 | red | red | T378 |
| 5237 | 324 | S | I | 0.003768 | 0.071438 | yes | 1.000000 | 1 | 1.000000 | 1 | red | red | S324 |
| 5239 | 324 | S | K | 0.003768 | -0.078653 | yes | 2.000000 | >1 | 2.000000 | >1 | gray | gray | S324 |

Also appending preferences of each amino acid in A549 cells for cross-referencing.

In [48]:

```
prefsdf = pd.melt(pd.read_csv('data/summary_avgprefs_rescaled.csv'),
        id_vars=['site'],
        value_vars=['A','C','D','E','F','G','H','I','K','L','M','N','P','Q','R','S','T','V','W','Y'],
        var_name='aa', value_name='pref'
       )
allmutdiffselmean = pd.merge(allmutdiffselmean, prefsdf, how='left',
                          left_on=['site', 'mutation'], right_on=['site', 'aa'])
allmutdiffselmean.head()
```

Out[48]:

|  | site | wildtype | mutation | mutdiffsel\_pim | mutdiffsel\_A549 | known | min Subst\_Avian | acc\_Avian | min Subst\_Human | acc\_Human | col\_Avian | col\_Human | site\_label | aa | pref |
| --- | --- | --- | --- | --- | --- | --- | --- | --- | --- | --- | --- | --- | --- | --- | --- |
| 0 | 431 | M | E | 6.299282 | -1.367148 | no | 2.000000 | >1 | 2.000000 | >1 | gray | gray | M431 | E | 0.080424 |
| 1 | 376 | K | A | 6.001965 | 0.834899 | no | 2.000000 | >1 | 2.000000 | >1 | gray | gray | K376 | A | 0.006534 |
| 2 | 357 | H | D | 5.569827 | 2.337165 | no | 1.000000 | 1 | 1.000000 | 1 | red | red | H357 | D | 0.301721 |
| 3 | 376 | K | R | 5.551795 | 2.105670 | yes | 0.999159 | 1 | 1.000000 | 1 | red | red | K376 | R | 0.118068 |
| 4 | 337 | S | H | 5.468044 | 1.152390 | no | 2.443822 | >1 | 2.998773 | >1 | gray | gray | S337 | H | 0.053208 |

##### A broad look at trends and top mutations¶

I want to distinguish sites responsible for non-specifically improving growth in A549 cells in the presence of pimodivir, versus those which are specific to pimodivir. To do so, I scatterplot the site differential selection values for selection in 50nM pimodivir, vs selection in human A549 vs duck CCL-141 cells.

In [60]:

```
%matplotlib inline
with sns.axes_style("darkgrid"):
    fig, axes = plt.subplots(1,1,figsize=(7,5))
    sns.scatterplot('positive_diffsel_A549', 'positive_diffsel_pim',
                    hue='known', hue_order=['no', 'yes'],
                    alpha=0.5,
                   data=allsitediffselmean)
    plt.legend(bbox_to_anchor=(1.05, 1), loc=2, borderaxespad=0.)
    plt.tight_layout()
```

The sites in the top left quadrant represent sites having mutations that enhance replication specifically in the presence of pimodivir, but not improving replication in A549 over CCL141 cells.

In [61]:

```
pimodivirNOTa549_topsites = (allsitediffselmean.query(('positive_diffsel_pim > 8 & positive_diffsel_A549 < 5'))
                         .sort_values(by=['known', 'positive_diffsel_pim'], ascending=False)
                        )
# selectedsites = list(pimodivirNOTa549_topsites['site'])
display(HTML(pimodivirNOTa549_topsites.to_html(index=False)))
```

| site | abs\_diffsel\_pim | positive\_diffsel\_pim | abs\_diffsel\_A549 | positive\_diffsel\_A549 | known |
| --- | --- | --- | --- | --- | --- |
| 431 | 44.999962 | 44.934134 | 29.827025 | 0.518602 | yes |
| 337 | 40.598895 | 40.598895 | 10.303483 | 1.700791 | yes |
| 510 | 13.014941 | 13.014941 | 19.888240 | 1.380712 | yes |
| 363 | 11.268616 | 11.111315 | 8.613014 | 2.393958 | yes |
| 406 | 9.303846 | 8.805432 | 15.081346 | 0.098451 | yes |
| 196 | 23.368766 | 23.030144 | 18.903253 | 0.198914 | no |
| 302 | 21.014377 | 20.913276 | 19.311978 | 0.439029 | no |
| 677 | 22.618323 | 18.533366 | 7.769580 | 2.935992 | no |
| 188 | 18.586740 | 17.111561 | 9.856379 | 4.963079 | no |
| 439 | 17.085248 | 15.161841 | 11.942019 | 0.723882 | no |
| 195 | 17.428045 | 15.150804 | 13.319776 | 0.883846 | no |
| 256 | 14.914808 | 14.581982 | 10.013759 | 4.749988 | no |
| 611 | 17.088467 | 13.359814 | 9.855708 | 2.834170 | no |
| 360 | 14.638600 | 12.992224 | 38.990550 | 0.224438 | no |
| 74 | 13.762472 | 12.264116 | 9.530880 | 0.892474 | no |
| 300 | 11.977987 | 10.557809 | 12.285597 | 3.520387 | no |
| 377 | 13.014817 | 10.312068 | 38.073277 | 0.000000 | no |
| 491 | 11.386572 | 9.675054 | 20.338993 | 0.451335 | no |
| 333 | 9.599279 | 9.181033 | 9.364918 | 2.565604 | no |
| 342 | 12.396846 | 8.763625 | 16.299508 | 1.555916 | no |
| 511 | 9.272000 | 8.345721 | 16.387576 | 0.687005 | no |
| 464 | 10.531326 | 8.292640 | 16.623725 | 1.454258 | no |
| 320 | 9.402350 | 8.227207 | 16.010249 | 1.495925 | no |
| 400 | 8.676421 | 8.223686 | 8.468275 | 1.340710 | no |
| 153 | 8.351603 | 8.188177 | 6.197808 | 3.897498 | no |
| 149 | 8.346576 | 8.182106 | 5.828120 | 1.995725 | no |
| 472 | 11.910792 | 8.063840 | 17.521125 | 0.398978 | no |

The sites in the top right quadrant represent mutations that enhance replication specifically in the presence of pimodivir, but that also improve replication in A549 over CCL141 cells. Hence, they may be non-specifically improving replication.

In [62]:

```
pimodivirANDa549_topsites = (allsitediffselmean.query(('positive_diffsel_pim > 8 & positive_diffsel_A549 > 5'))
                         .sort_values(by='positive_diffsel_pim', ascending=False)
                        )
# selectedsites = list(pimodivirANDa549_topsites['site'])
display(HTML(pimodivirANDa549_topsites.to_html(index=False)))
```

| site | abs\_diffsel\_pim | positive\_diffsel\_pim | abs\_diffsel\_A549 | positive\_diffsel\_A549 | known |
| --- | --- | --- | --- | --- | --- |
| 357 | 41.789181 | 41.789181 | 17.474837 | 14.032147 | yes |
| 323 | 39.914311 | 39.876937 | 21.051950 | 19.962100 | yes |
| 376 | 35.435920 | 35.394002 | 12.082695 | 11.358961 | yes |
| 627 | 35.990894 | 33.737983 | 19.589407 | 18.009501 | no |
| 514 | 27.157629 | 27.157629 | 10.452031 | 5.027741 | no |
| 192 | 21.268584 | 21.257035 | 9.903482 | 6.898806 | no |
| 701 | 16.009550 | 15.563750 | 18.423070 | 17.946994 | no |
| 304 | 16.314875 | 15.126318 | 18.501685 | 7.079232 | no |
| 303 | 13.561978 | 11.417506 | 15.880285 | 7.834892 | no |
| 588 | 14.832157 | 11.245627 | 12.154298 | 9.662929 | no |
| 185 | 12.040127 | 9.587045 | 10.449278 | 6.749344 | no |
| 158 | 11.557029 | 9.575199 | 13.200199 | 12.652284 | no |
| 521 | 13.264346 | 8.920490 | 34.523299 | 31.595969 | no |
| 715 | 10.872807 | 8.546653 | 9.744296 | 5.008080 | no |

I also scatterplot differential selection values for each mutation under selection in 50nM pimodivir, vs selection in human A549 vs duck CCL-141 cells. Known mutations have positive mutdiffsel in pimodivir, but have a range of mutdiffsel as measured under selection in human vs duck cells.

We also see that most pimodivir resistance mutations don't have high preference. I suspect this is because mutations are tolerated at sites that are tolerant of many mutations, so the preference for any one mutation will be low.

In [63]:

```
with sns.axes_style("darkgrid"):
    fig, axes = plt.subplots(1,2,figsize=(12,5))
    sns.scatterplot('mutdiffsel_A549', 'mutdiffsel_pim',
                    hue='known', hue_order=['no', 'yes'],
                    alpha=0.5,ax=axes[0],
                   data=allmutdiffselmean)
    sns.scatterplot('pref', 'mutdiffsel_pim',
                    hue='known', hue_order=['no', 'yes'],
                    alpha=0.5,ax=axes[1],
                   data=allmutdiffselmean)
    plt.legend(bbox_to_anchor=(1.05, 1), loc=2, borderaxespad=0.)
    plt.tight_layout()
```

Printing some of the most highly differentially selected mutations to look at.

In [64]:

```
mutdiffselmean_top = allmutdiffselmean.query(('mutdiffsel_pim > 4'))
selectedmuts = list(mutdiffselmean_top['site'])
display(HTML(mutdiffselmean_top.to_html(index=False)))
```

| site | wildtype | mutation | mutdiffsel\_pim | mutdiffsel\_A549 | known | min Subst\_Avian | acc\_Avian | min Subst\_Human | acc\_Human | col\_Avian | col\_Human | site\_label | aa | pref |
| --- | --- | --- | --- | --- | --- | --- | --- | --- | --- | --- | --- | --- | --- | --- |
| 431 | M | E | 6.299282 | -1.367148 | no | 2.000000 | >1 | 2.000000 | >1 | gray | gray | M431 | E | 0.080424 |
| 376 | K | A | 6.001965 | 0.834899 | no | 2.000000 | >1 | 2.000000 | >1 | gray | gray | K376 | A | 0.006534 |
| 357 | H | D | 5.569827 | 2.337165 | no | 1.000000 | 1 | 1.000000 | 1 | red | red | H357 | D | 0.301721 |
| 376 | K | R | 5.551795 | 2.105670 | yes | 0.999159 | 1 | 1.000000 | 1 | red | red | K376 | R | 0.118068 |
| 337 | S | H | 5.468044 | 1.152390 | no | 2.443822 | >1 | 2.998773 | >1 | gray | gray | S337 | H | 0.053208 |
| 677 | E | L | 5.416295 | -0.178586 | no | 1.999720 | >1 | 2.000000 | >1 | gray | gray | E677 | L | 0.024065 |
| 514 | S | R | 5.367328 | 0.954636 | no | 2.000000 | >1 | 2.000000 | >1 | gray | gray | S514 | R | 0.080986 |
| 323 | F | T | 5.364631 | 1.333052 | no | 2.000000 | >1 | 2.000000 | >1 | gray | gray | F323 | T | 0.030985 |
| 431 | M | Q | 5.307767 | -0.514680 | no | 2.000000 | >1 | 2.000000 | >1 | gray | gray | M431 | Q | 0.019455 |
| 376 | K | Q | 5.209439 | 0.804407 | no | 1.000841 | 1 | 1.000000 | 1 | red | red | K376 | Q | 0.018076 |
| 337 | S | A | 5.052127 | 0.013794 | no | 1.000000 | 1 | 1.000000 | 1 | red | red | S337 | A | 0.049033 |
| 431 | M | H | 4.942171 | -1.737010 | no | 3.000000 | >1 | 3.000000 | >1 | gray | gray | M431 | H | 0.016466 |
| 337 | S | C | 4.853308 | -0.474358 | no | 1.444102 | >1 | 1.998773 | >1 | gray | gray | S337 | C | 0.041835 |
| 323 | F | L | 4.776464 | 3.174165 | no | 0.999440 | 1 | 1.000000 | 1 | red | red | F323 | L | 0.200610 |
| 363 | F | L | 4.729033 | -1.375224 | no | 0.999159 | 1 | 1.000000 | 1 | red | red | F363 | L | 0.006302 |
| 323 | F | I | 4.706787 | 2.079226 | no | 1.000000 | 1 | 1.000000 | 1 | red | red | F323 | I | 0.104682 |
| 514 | S | Q | 4.618401 | -0.866545 | no | 2.555338 | >1 | 2.999755 | >1 | gray | gray | S514 | Q | 0.198142 |
| 323 | F | C | 4.547562 | 1.301712 | no | 1.000560 | 1 | 1.000000 | 1 | red | red | F323 | C | 0.088432 |
| 431 | M | G | 4.479676 | -1.601653 | no | 2.000000 | >1 | 2.000000 | >1 | gray | gray | M431 | G | 0.009253 |
| 376 | K | S | 4.478762 | 0.728452 | no | 1.999159 | >1 | 2.000000 | >1 | gray | gray | K376 | S | 0.009541 |
| 337 | S | G | 4.442308 | -0.152359 | no | 2.000000 | >1 | 2.000000 | >1 | gray | gray | S337 | G | 0.017084 |
| 357 | H | L | 4.440923 | 1.718452 | no | 1.001961 | 1 | 1.000000 | 1 | red | red | H357 | L | 0.008802 |
| 431 | M | P | 4.219925 | 0.518602 | no | 2.000000 | >1 | 2.000000 | >1 | gray | gray | M431 | P | 0.040982 |
| 192 | E | R | 4.189553 | -0.713758 | no | 2.000000 | >1 | 2.000000 | >1 | gray | gray | E192 | R | 0.019884 |
| 188 | E | R | 4.134855 | 0.475821 | no | 1.995797 | >1 | 1.998527 | >1 | gray | gray | E188 | R | 0.020751 |
| 323 | F | V | 4.132682 | 2.109604 | no | 1.000000 | 1 | 1.000000 | 1 | red | red | F323 | V | 0.183100 |
| 149 | P | V | 4.037609 | 0.414441 | no | 2.000000 | >1 | 2.000000 | >1 | gray | gray | P149 | V | 0.028434 |
| 302 | P | S | 4.029952 | -1.008011 | no | 1.000280 | 1 | 1.000000 | 1 | red | red | P302 | S | 0.021016 |
| 627 | E | L | 4.013941 | 0.856598 | no | 1.995517 | >1 | 2.000000 | >1 | gray | gray | E627 | L | 0.031988 |

##### Plotting individual replicates to aid selection of individual mutations for validation¶

I want to select mutations that are reproducible selection across replicates. Hence, I collate data from all replicates for plotting.

In [65]:

```
selection = 'Pimodivir50nM'
reps = [1,2,3]
alldiffsel = {}
diffsellist = []
for rep in reps:
    diffsellist.append(((pd.read_csv(os.path.join(resultsdir['diffsel'], f"{selection}-rep{rep}_mutdiffsel.csv"))
                        .pivot(index='site', columns='mutation', values='mutdiffsel') )
                        .reset_index() )
                        .assign(replicate=rep, replicatelabel=f"rep{rep}")
                        )
alldiffsel[selection] = pd.concat(diffsellist)

def plotDiffselReplicates(selectedsites):
    fig, axes = plt.subplots(1,len(selectedsites),
                            figsize=(len(selectedsites)*2,3))
    for (i,site) in enumerate(selectedsites):
        alldiffselatsite = (alldiffsel[selection][(alldiffsel[selection]['site']==site)]
                            .melt(id_vars=['replicate','replicatelabel','site'],value_vars=dms_tools2.AAS,
                                  var_name='mutation', value_name='mutdiffsel')
                            .query('mutdiffsel > 0')
                            .sort_values(['replicate'])
                            )
        _ = dmslogo.draw_logo(data=alldiffselatsite,
                              x_col='replicate',
                              letter_col='mutation',
                              letter_height_col='mutdiffsel',
                              xtick_col='replicatelabel',
                              xlabel=f"site {site}",
                              ax=axes[i],
                              ylabel='mutdiffsel' if i==0 else ''
                             )
    plt.tight_layout()
```

Plot the sites with highest positive differential selection.

In [66]:

```
pimodivirNOTa549_topsites.sort_values(by="site", inplace=True)
plotDiffselReplicates((pimodivirNOTa549_topsites['site']).tolist()[0:9])
plotDiffselReplicates((pimodivirNOTa549_topsites['site']).tolist()[9:18])
plotDiffselReplicates((pimodivirNOTa549_topsites['site']).tolist()[18:])
```

I also plot here the sites that may have been selected due to host selection.

In [67]:

```
plotDiffselReplicates((pimodivirANDa549_topsites['site']).tolist())
```

Reassuringly, pimodivir resistance mutations have previously been identified at a number of these sites.

From Byrn et al., 2015, and in **bold** the sites that fall within the top sites plotted here: "Six PB2 variants, Q306H, S324I, S324N, S324R, F404Y, and **N510T**, were identified in two or more independent cultures and showed a greater than 10-fold change in VX-787 sensitivity."

From Finberg et al., 2019, and in **bold** the sites that fall within the top ten plotted here: "Analysis of samples collected after baseline while undergoing treatment demonstrated emergence of PB2 substitutions or phenotypic resistance to pimodivir in 11 patients (4, 6, and 1 in the pimodivir 300 mg, pimodivir 600 mg, and combination groups, respectively). The mutations included S324K/N/R, F325L, **S337P**, K376N/R, T378S, and **N510K**, and their emergence was typically associated with a concomitant increase in the pimodivir fold change (range, 9.4 to >372.0). Further details will be reported elsewhere."

Finberg et al., 2019 also note: "The following PB2 positions of interest (ie, the positions potentially associated with resistance to pimodivir) were defined on the basis of in vitro and human challenge study data: Q306, F323, S324, F325, **S337**, H357, F363, **K376**, T378, F404, Q406, **M431**, and **N510**." However, they do not identify the specific resistance mutations for these sites.

What do the other sites reported in Finberg (2019), but not within the top sites I plotted above, look like? I plot these below. We do see selection, often for the mutation reported in the paper. However they did not pass the above stringent thresholds.

In [68]:

```
selectedsites = [306, 323, 324, 325, 357, 363, 376, 378, 404, 406]
plotDiffselReplicates(selectedsites)
```

### Final selection of mutations to validate¶

I selected the following mutations to validate, through a combination of these data, information from the literature, and examining the mutations on the structures of PB2.

Make the following mutations on the S009-PB2-627K background, definitely on the HDM (1563), and maybe also pHW (1516) backbones.

Full list of mutations here. (Details and individual plots below.)

| Selected amino acid mutation to validate | Codon mutation to make | Notes |
| --- | --- | --- |
| F323L | TTC > CTT | We already previously made this mutation on the HDM backbone (2537:HDM\_S009\_PB2-F323L-E627K), but not the pHW version yet. |
| H357D | CAT > GAC |  |
| D256G | GAC > GGT |  |
| P302S | CCA > CTC |  |
| M431E | ATC > GAA |  |
| S514R | TCA > CGT |  |
| E192R | GAA > GCT |  |
| C196H | TGT > CAC |  |

In [69]:

```
sites_direct = [323, 337] + list(range(355, 364)) + list(range(376, 379)) + list(range(404, 407))
sites_apo = [256] + list(range(300, 307)) + list(range(324, 328)) + [431, 432] + list(range(510, 518))
sites_nterm = list(range(186, 199))
sites_human = [627] + list(range(674, 678)) + [701] # Other/human adaptive
```

In [70]:

```
def linelogoprefplot(sites_to_show):
    df = (allmutdiffselmean
          .assign(site_label=lambda x: x['wildtype'] + x['site'].map(str),
                  show_site=lambda x: x['site'].isin(sites_to_show),
                 )
          .query('show_site')
         )
    dfsite = allsitediffselmean.assign(show_site=lambda x: x['site'].isin(sites_to_show),)

    fig, axes = plt.subplots(3, 1)
    fig.subplots_adjust(hspace=0.3) # add more vertical space for axis titles
    fig.set_size_inches(len(sites_to_show), 16)

    # draw diffsel plot, no x-axis ticks or label, default coloring
    _ = dmslogo.draw_logo(df.assign(no_ticks=''),
                          x_col='site',
                          letter_col='mutation',
                          letter_height_col='mutdiffsel_pim',
                          ax=axes[0],
                          xlabel='',
                          ylabel='',
                          xtick_col='no_ticks',
                          clip_negative_heights=True,
                          title='')

    # draw diffsel plot, color as specified in `data`
    _ = dmslogo.draw_logo(df.assign(no_ticks=''),
                          x_col='site',
                          letter_col='mutation',
                          letter_height_col='mutdiffsel_pim',
                          color_col='col_Avian',
                          ax=axes[1],
                          xlabel='',
                          ylabel='',
                          xtick_col='no_ticks',
                          clip_negative_heights=True,
                          title='')
    
    # draw pref plot, default coloring
    _ = dmslogo.draw_logo(df,
                          x_col='site',
                          letter_col='mutation',
                          letter_height_col='pref',
                          ax=axes[2],
                          xlabel='',
                          ylabel='',
                          xtick_col='site_label',
                          clip_negative_heights=True,
                          title='')

    fig, ax = dmslogo.draw_line(dfsite,
                                x_col='site',
                                height_col='positive_diffsel_pim',
                                xtick_col='site',
                                show_col='show_site',
                                title='',
                                widthscale=2.5)
```

#### Sites of resistance mutations likely to be in close contact with m7G or pimodivir¶

| Selected amino acid mutation to validate | Codon mutation to make |
| --- | --- |
| F323L | TTC > CTT |
| H357D | CAT > GAC |

In [71]:

```
linelogoprefplot(sites_direct)
```

#### Sites of resistance mutations in mid-link domain, likely to stabilize apo form of polymerase¶

| Selected amino acid mutation to validate | Codon mutation to make |
| --- | --- |
| D256G | GAC > GGT |
| P302S | CCA > CTC |
| M431E | ATC > GAA |
| S514R | TCA > CGT |

In [72]:

```
linelogoprefplot(sites_apo)
```

#### Sites of resistance mutations in N-terminal domain of PB2¶

| Selected amino acid mutation to validate | Codon mutation to make |
| --- | --- |
| E192R | GAA > GCT |
| C196H | TGT > CAC |

In [73]:

```
linelogoprefplot(sites_nterm)
```

#### Sites of resistance mutations that I found in my previous screen to be human adaptive, or others¶

Just plotting, not validating any of these.

In [74]:

```
linelogoprefplot(sites_human)
```

In [ ]:
